# Universal gene co-expression network reveals receptor-like protein genes conferring broad-spectrum resistance in pepper (*Capsicum annuum* L.)

**DOI:** 10.1101/2021.03.04.433825

**Authors:** Won-Hee Kang, Junesung Lee, Namjin Koo, Ji-Su Kwon, Boseul Park, Yong-Min Kim, Seon-In Yeom

**Author notes:** Corresponding author: Seon-In Yeom, Won-Hee Kang.

## Abstract

Receptor-like proteins (RLPs) on the plant cell surface have been implicated in immune responses and developmental processes. Although hundreds of *RLP* genes have been identified in plants, only a few RLPs have been functionally characterized in a limited number of plant species. Here, we identified *RLPs* in the pepper (*Capsicum annuum*) genome, and performed comparative transcriptomics coupled with the analysis of conserved gene co-expression networks (GCNs) to reveal the role of core RLP regulators in pepper–pathogen interactions. A total of 102 RNA-seq datasets of pepper plants infected with four pathogens were used to construct CaRLP-targeted GCNs (CaRLP-GCN). All resistance-responsive CaRLP-GCNs were merged to construct a universal GCN. Fourteen hub *CaRLPs*, tightly connected with defense related gene clusters, were identified in eight modules. Based on the CaRLP-GCNs, we experimentally tested whether hub *CaRLPs* in the universal GCN are involved in biotic stress response. Of the nine hub *CaRLPs* tested by virus-induced gene silencing (VIGS), three genes (*CaRLP264*, *CaRLP277*, and *CaRLP351*) showed defense suppression with less hypersensitive response (HR)-like cell death in race-specific and non-host resistance response to viruses and bacteria, respectively, and consistently enhanced susceptibility to *Ralstonia solanacearum* and/or *Phytophthora capsici*. These data suggest that key *CaRLPs* exhibit conserved functions in response to multiple biotic stresses and can be used for engineering of a plant with broad-spectrum resistance. Altogether, we show that generation of a universal GCN using comprehensive transcriptome datasets could provide important clues for uncovering genes involved in various biological processes.

## INTRODUCTION

Plants employ extra- and intracellular immune signaling to protect themselves against pathogens^1,2^. The first layer of plant immunity, known as pattern-triggered immunity, is activated upon the perception of pathogen- or microbe-associated molecular patterns (PAMPs or MAMPs) by plant cell surface-localized pattern recognition receptors (PRRs). PRRs sense diverse pathogens including bacteria, fungi, oomycetes and parasitic plants, and are involved in the immune signaling complex and network^3^. Recently, plant PRRs have been successfully used to confer broad-spectrum resistance in potato (*Solanum tuberosum*)^4,5^ and in *Nicotiana benthamiana* and tomato (*Solanum lycopersicum*) (Lacombe et al., 2010), and have been considered for conferring broad-spectrum disease resistance in other crops.

Plant PRRs are distinguished into two main classes, depending on their cytoplasmic kinase domains: receptor-like kinases (RLKs) and receptor-like proteins (RLPs). RLKs contain an extracellular domain, a single transmembrane domain and a cytoplasmic domain, whereas RLPs lack the cytoplasmic kinase domain but carry a short cytoplasmic tail. RLPs play crucial roles in plant immunity against pathogens. The first RLPs, designated as *Cf* genes, were identified in tomato, which imparted resistance to *Cladosporium fulvum* isolates^6–9^. Since then, several RLPs have been shown to have function in plant defense, mostly in Solanaceous plants and *Arabidopsis thaliana*^5,10–18^. In addition, RLPs are also involved in plant development ^19–21^. A number of genes encoding RLPs have been identified with the completion of plant genome project^22–27^; however, relatively fewer genes have been functionally characterized to date.

Based on the recent advances in the sequencing technology, along with the decline in the cost of sequencing, RNA-seq have been widely utilized in plants, producing massive amounts of data. However, the identification and manipulation of information of interest from large integrated datasets remain challenging. Since functionally associated genes often show transcriptional co-regulation, gene co-expression networks (GCNs) present an important resource for the identification of novel genes within a given biological process-regulating module. Thus, the analysis of GCNs could be a powerful approach for predicting gene functions and isolating modules involved in specific biological process across large-scale gene expression data^28–31^. In recent years, GCN analysis has been successfully used to discover stress-responsive genes in plants^32–34^. Additionally, several research groups recently performed comparative and combined analyses of GCNs in time-series experiments conducted under various conditions and with multiple treatments, across different species and kingdoms^35–39^. These studies were used to identify hub genes and infer their roles in biological processes. However, that are less well investigated compared with certain model species because of their extreme complexity and limited resources.

Chili pepper (*Capsicum* spp.), a member of the Solanaceae family, is an important vegetable crop worldwide. However, pepper production is threatened by pathogens such as fungi, bacteria, viruses, insects, and nematodes. Development of pathogen resistant cultivars is one of the best approaches for controlling infection in pepper. Although multiple reference genomes and transcriptome datasets of pepper have been published recently^40–43^, the molecular mechanism underlying plant immunity remains unclear. Therefore, the identification and characterization of genes involved in plant defense using comprehensive transcriptome data is critical. In this study, we identified 438 RLP genes in the chili pepper genome through phylogenetic analysis and comparative transcriptomic analysis of 102 RNA-seq datasets of chili pepper plants challenged with four different pathogens. In addition, we constructed CaRLP-targeted GCNs (CaRLP-GCN) using comprehensive RNA-seq datasets and, merged the resistant-responsive GCNs to develop a universal CaRLP-GCN. Using this GCN, we identified 14 putative RLP hub genes belonging to eight modules. Loss-of-function analysis of three *CaRLPs* (*CaRLP264*, *CaRLP277*, and *CaRLP351*) validated the broad immune response to pathogens. The silencing of each of the three *CaRLPs* significantly reduced the broad-spectrum resistance against viruses, bacteria, and oomycetes. Overall, this study demonstrates the successful characterization of novel genes via the construction of a universal GCN from large RNA-seq datasets, and provides key insights into the broad-spectrum resistance in plants.

## RESULTS

### Genome-wide identification and classification of RLPs in pepper genome

A total of 438 RLP-encoding genes were identified in the *Capsicum annuum* genome by excluding redundant sequences and genes encoding NB-ARC or kinase domain-containing proteins, and by validating the RLP structure (see Materials and Methods for details). All of the RLP-encoding genes were renamed according to their chromosomal positions (Fig. 1a and Supplementary Table S1). Details of *CaRLP* genes are summarized in Supplementary Table S1.

**Fig. 1.**
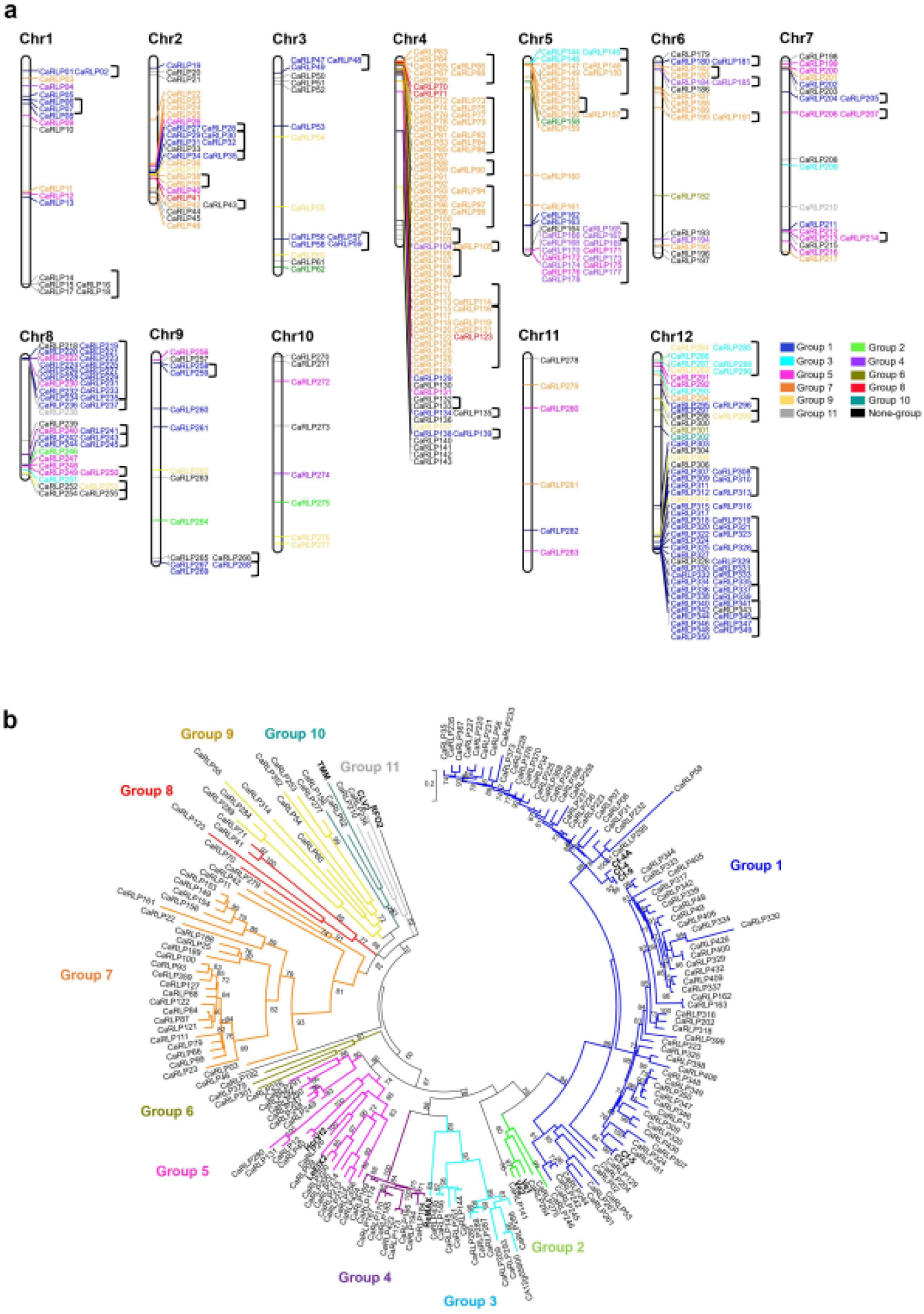
Phylogenetic analysis and chromosomal locations of *CaRLPs*. (a) Physical location of *CaRLP* genes on pepper chromosomes. A total of 350 *CaRLPs* were located on 12 chromosomes, while 88 *CaRLPs* were assigned to pepper scaffolds. The chromosome and scaffold numbers are indicated at the top of each chromosome. *CaRLPs* are colored according to their phylogenetic group. Black square brackets on the right side of gene IDs indicate the physical gene clusters of CaRLPs on chromosomes. (b) Phylogenetic tree of CaRLPs constructed using the maximum-likelihood method using PhyML. Bootstrap values over 60 are indicated above branches. Clades containing known *RLP* genes are indicated by the *RLP* gene names at the end of the clade.

Phylogenetic analysis and sequence similarity-based clustering methods^24,44^ classified 364 out of 438 CaRLPs into 11 groups; the remaining 74 CaRLPs were not classified into any group and were defined as singletons (Fig. 1b and Supplementary Table S1). The majority of RLPs were assigned to two groups (1 and 7). Group 1 was the largest group consisting of comprising 153 genes, including tomato *SlCf* genes and their homologs in pepper. Group 7 was the second largest group comprising 118 *CaRLP* genes, without known genes. Next, to explore the evolutionary relationships of CaRLPs with SlRLPs^24^ and AtRLPs^22^, we conducted phylogenetic analysis of the amino acid sequence of conserved C3-D domain of these RLPs (Supplementary Fig. S1). The majority of CaRLPs clustered together with SlRLPs, whereas most of the AtRLPs grouped separately, forming two Arabidopsis-specific clades (Supplementary Fig. S1). These results suggest that the *CaRLP* gene family underwent expansion after divergence from the common ancestor of Arabidopsis and Solanaceae species.

### Chromosomal location, physical cluster, and conserved motif analyses

Of the 438 *CaRLP* genes identified in this study, 350 mapped on to 12 chromosomes, while 88 were assigned to unmapped scaffolds (Fig. 1a). Most of the *CaRLPs* belonging to the same phylogenetic group were closely clustered on a given chromosome. Next, we performed physical cluster analysis to thoroughly investigate the chromosomal distribution of *CaRLPs*. The results revealed 54 clusters on pepper chromosomes containing 227 genes (Fig. 1a and Supplementary Table S2). Each of these clusters spanned a physical distance of 0.7–885 kb. Large clusters (>200 kb) were located on chromosomes 1, 4, 5, 8, and 12, and no cluster was identified on chromosomes 10 and 11. Of all the *CaRLPs* in each group, those with large numbers (Groups 1 and 7) formed mostly physical clusters. To better understand the *CaRLP* gene family, we examined conserved motifs in CaRLP proteins. A total of 20 distinct motifs were predicted among all 438 CaRLPs and known RLPs (Fig. S2 and Supplementary Table S3). Most motifs were found to encode the leucine-rich repeat (LRR) domain, while motif 9 encoded the transmembrane region. Most of the closely related genes in the same phylogenetic group exhibited common motif compositions. Taken together, these data indicate that CaRLPs belonging to the same phylogenetic group share conserved motifs, similar protein domain compositions and similar chromosomal locations.

### Expression analysis of *CaRLPs* in response to biotic stresses

RLPs perform crucial roles in plant disease resistance. However, little is known about the possible function of CaRLPs in defense response. To further understand the role of *CaRLP* genes in plant defense, we investigated the expression patterns of *CaRLPs* showing differential expression between uninoculated (control) and pathogen-challenged pepper plants; these genes are hereafter referred to as differentially expressed *CaRLP* genes (*CaRLP*-DEGs). These DEGs were obtained from 63 previously published RNA-seq datasets of pepper plants infected with three different viruses including *Tobacco mosaic virus* (TMV) pathotype P0 (TMV-P0), TMV pathotype P2 (TMV-P2) and *Pepper mottle virus* (PepMoV)^43,45^. In addition, to determine the changes in *CaRLP* expression at an early stage of oomycete infection, we generated six-timepoint RNA-seq datasets from three biological replicates of *P. capsici*-inoculated and control pepper plants (Supplementary Table S4). Thus, we examined a total of 102 RNA-seq datasets to analyze the expression of *CaRLPs* (Supplementary Table S5). Pepper accession ‘CM334,’ which was used for RNA-seq analysis in this study, is known to be resistant to TMV-P0, PepMoV, and *P. capsici* but susceptible to TMV-P2^45,46^.

Of the 438 *CaRLPs*, 35 were differentially expressed between TMV-P0-inoculated and control plants, and 6 were differentially expressed between PepMoV-inoculated and control plants (fold-change ≥ 2) at one or more time points (Supplementary Fig. S3, and Supplementary Table S6). However, no *CaRLP*-DEG was identified between TMV-P2-inoculated and control plants, susceptible response (Supplementary Fig. S3). Heat map analysis divided the identified DEGs into four hierarchical clusters (Fig. 2a and 2b). In each cluster, *CaRLP*-DEGs identified between TMV-P0 vs. control treatments showed dynamic expression patterns, unlike those identified in TMV-P2 vs. control and PepMoV vs. control treatments. Cluster 1 was enriched in *CaRLPs* down-regulated in TMV-P0- and PepMoV-infected plants at 72 h post-inoculation (hpi). *CaRLP*-DEGs in clusters 2, 3, and 4 were up-regulation at later time points, mainly in TMV-P0-inoculated plants. These results indicate that several *CaRLPs* are involved in the response to viral pathogens.

**Fig. 2.**
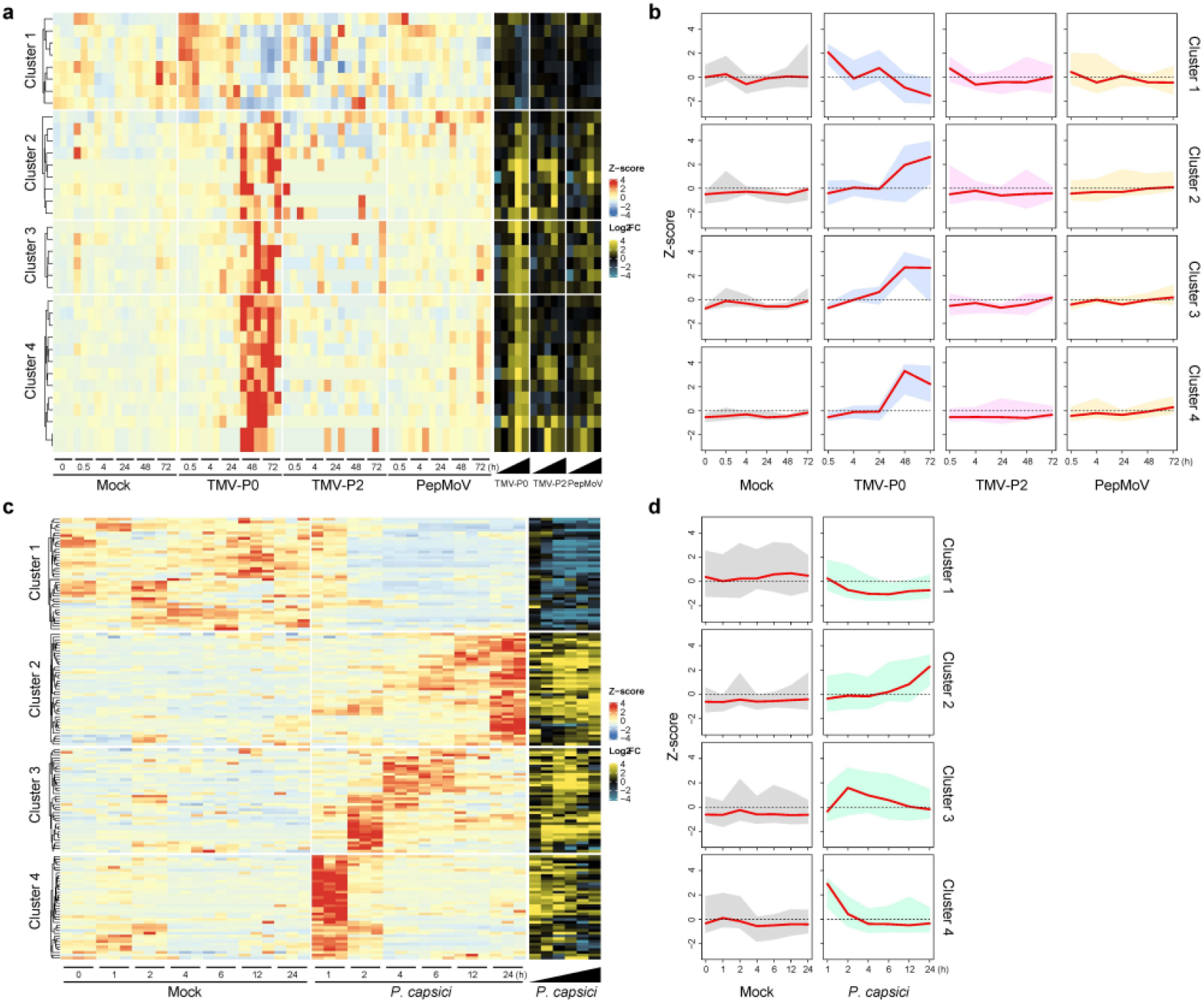
Analysis of *CaRLP* expression patterns during pathogen infection. (a) Heat map displaying the time-course expression profiles of differentially expressed genes (DEGs) identified in plants treated with TMV-P0, TMV-P2, and PepMoV. The left hand side (red to blue scale) and right hand side (yellow to blue scale) of the heatmap indicate DEGs with z-score and log_2_(fold-change; FC) values, respectively. (b) Time-course analysis of the expression pattern of *CaRLP-*DEGs within clusters. Each cluster represents the hierarchical clustering numbers in the heatmap shown in (a). Numbers represent the mean z-score of DEGs, and red lines indicate median z-score within a cluster. (c) Heat map illustrating the time-course expression profiles of *CaRLP*-DEGs identified in plants treated with *P. capsici*. (d) Expression profiling of CaRLP-DEGs in clusters under *P. capsici* infection.

Next, we compared the transcriptome of *P. capsici*-inoculated plants at 1, 2, 4, 6, 12, and 24 hpi with that of control plants, and identified 158 *CaRLP*-DEGs (fold-change ≥ 2) at one or more time points (Fig. 2c, 2d, and Supplementary Table S7). This number was much higher than that obtained from virus-inoculated plants. These 158 *CaRLPs* were also divided into four clusters by hierarchical clustering analysis. *CaRLP*-DEGs overrepresented in clusters 1 and 4 were down-regulated, whereas those in cluster 2 were up-regulated at later time points. Genes in cluster 3 were highly expressed at 1 hpi, but their expression decreased over time. A total of 31 *CaRLPs* were identified in both virus- and *P. capsici*-inoculated plants, and were referred to as common *CaRLP*-DEGs (Supplementary Table S6 and S7). Most of these 31 *CaRLP*-DEGs showed an increase in expression over time in both virus- and *P. capsici*-inoculated plants, and were classified into clusters 2, 3, and 4 in virus RNA-seq data and into clusters 2 and 3 in *P. capsici* RNA-seq data (Fig. 2). Taken together, comprehensive transcriptome profiling supported that CaRLPs are involved in an immune response against biotic stresses including viruses and oomycetes.

### Construction of comprehensive co-expression networks of *CaRLPs* using RNA-seq data of pathogen-challenged pepper plants

To understand functional implications of *CaRLPs* expressed during pathogen infection, CaRLP-targeted GCNs were constructed using all 102 RNA-seq datasets (described above). Four GCNs involving CaRLPs as hub genes were identified, one each from the RNA-seq data of TMV-P0-, TMV-P2-, PepMoV-, and *P. capsici*-challenged plants; these CaRLP-GCNs consisted of 4,041 nodes with 11,825 edges, 1,073 nodes with 1,194 edges, 3,732 nodes with 7,933 edges and 10,878 nodes with 84,255 edges, respectively (Fig. 3a, 3b, 3c and 3d). Gene ontology (GO) enrichment analysis was performed for the modules in each of the GCNs identified. Various GO terms were enriched in the molecular function (MF), cellular component (CC) and biological process (BP) categories. Interestingly, two GO terms, “oxidation-reduction process” and “cellular oxidation detoxification,” were overrepresented in the BP category in the three pathogen treated datasets of ‘CM334,’ which showed resistance response to TMV-P0, PepMoV, and *P. capsici* (Fig. 3e). These two biological processes are known to be involved in plant immune response: reduction-oxidation changes occur in response to pathogen invasion and subsequently activate the plant immune function, i.e., HR, a programmed execution of challenged plant cells ^47^; cellular oxidation detoxification has also been reported in plants under stress ^48^. The GO term “phosphorylation” was enriched in the GCN from TMV-P2 treated RNA-seq dataset. Previously, ^49^ reported that phosphorylation is induced upon plant virus infection. These findings suggest that CaRLPs and the corresponding genes in GCNs are involved in biotic stress response in pepper. In addition, we carried out Kyoto Encyclopedia of Genes and Genomes (KEGG) pathway analysis of genes belonging to each GCN. The results showed enrichment of pathways associated with plant immune response such as “biosynthesis of antibiotics,” “phenylalanine metabolism” and “phenylpropanoid biosynthesis” in TMV-P0-, PepMoV-, and *P. capsici*-specific CaRLP-GCNs, respectively (Supplementary Fig. S4). Taken together, GO and KEGG enrichment analyses of GCNs showed that genes connected with CaRLPs in GCNs are potentially involved in immune response in pepper.

**Fig. 3.**
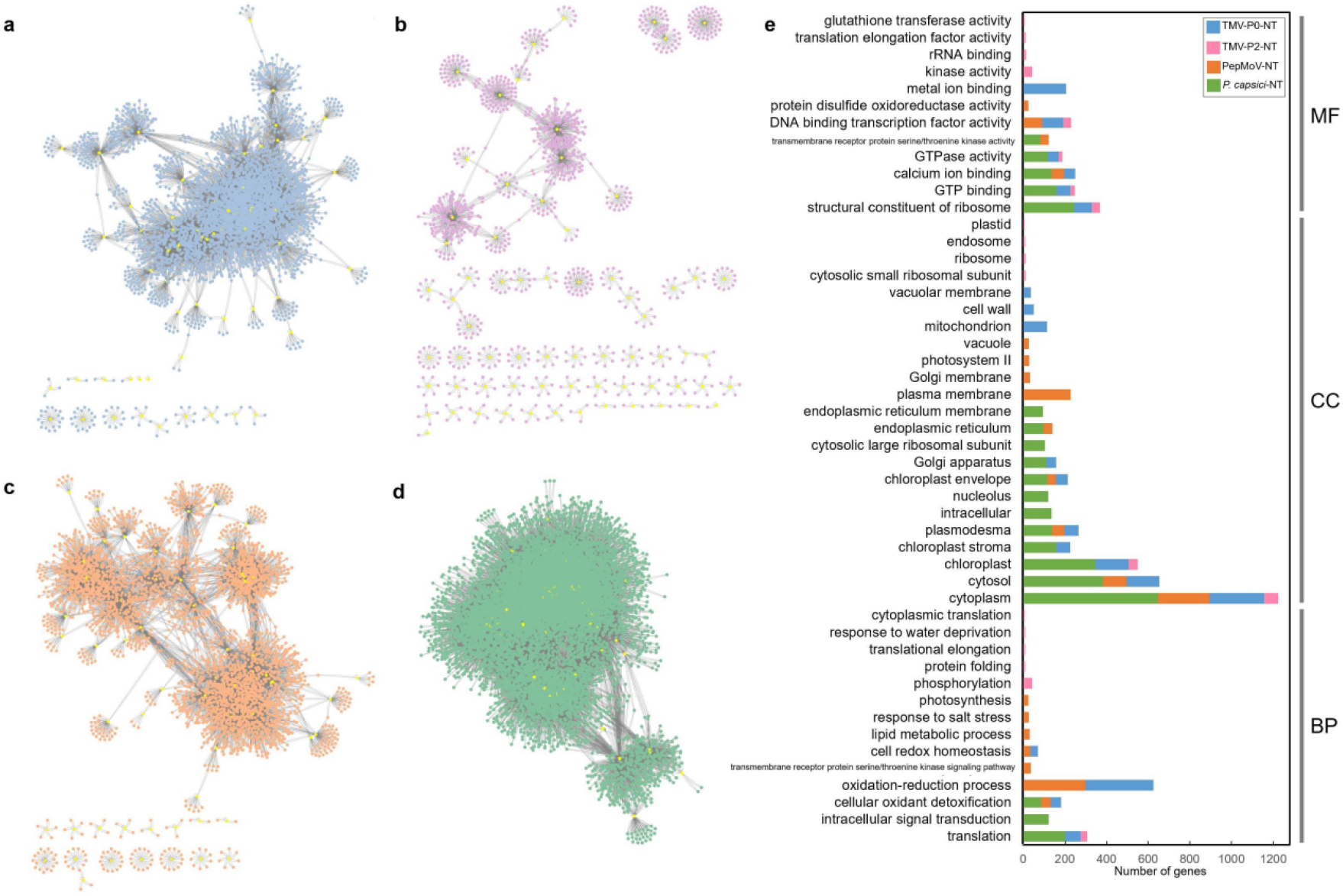
Analysis of the co-expression network (GCN) of *CaRLPs* identified using RNA-seq data of biotic stress treated pepper plants. (a–d) GCN comprising *CaRLP* hub genes identified in plants treated with TMV-P0 (a), TMV-P2 (b), PepMoV (c) and *P. capsici* (d). Yellow dots in the GCN indicate *CaRLPs*. (e) Top 20 GO terms significantly enriched in each of the four GCNs. The top 20 GO terms (*P* < 0.01) were selected in order of the largest number of genes from each of the four GCNs. BP, biological process; CC, cellular component; MF, molecular function.

### Identification of biotic stress-responsive core *CaRLPs*

To identify *CaRLPs* confer resistance to multiple pathogens, we merged the CaRLP-GCNs derived from the RNA-seq datasets of TMV-P0-, PepMoV-, and *P. capsici*-infected plants, thus constructing a universal resistance-responsive GCN (hereafter referred to as RN). The RN contained eight modules (named as RN1–8), with a total 14 hub *CaRLPs* (Fig. 4a).

**Fig. 4.**
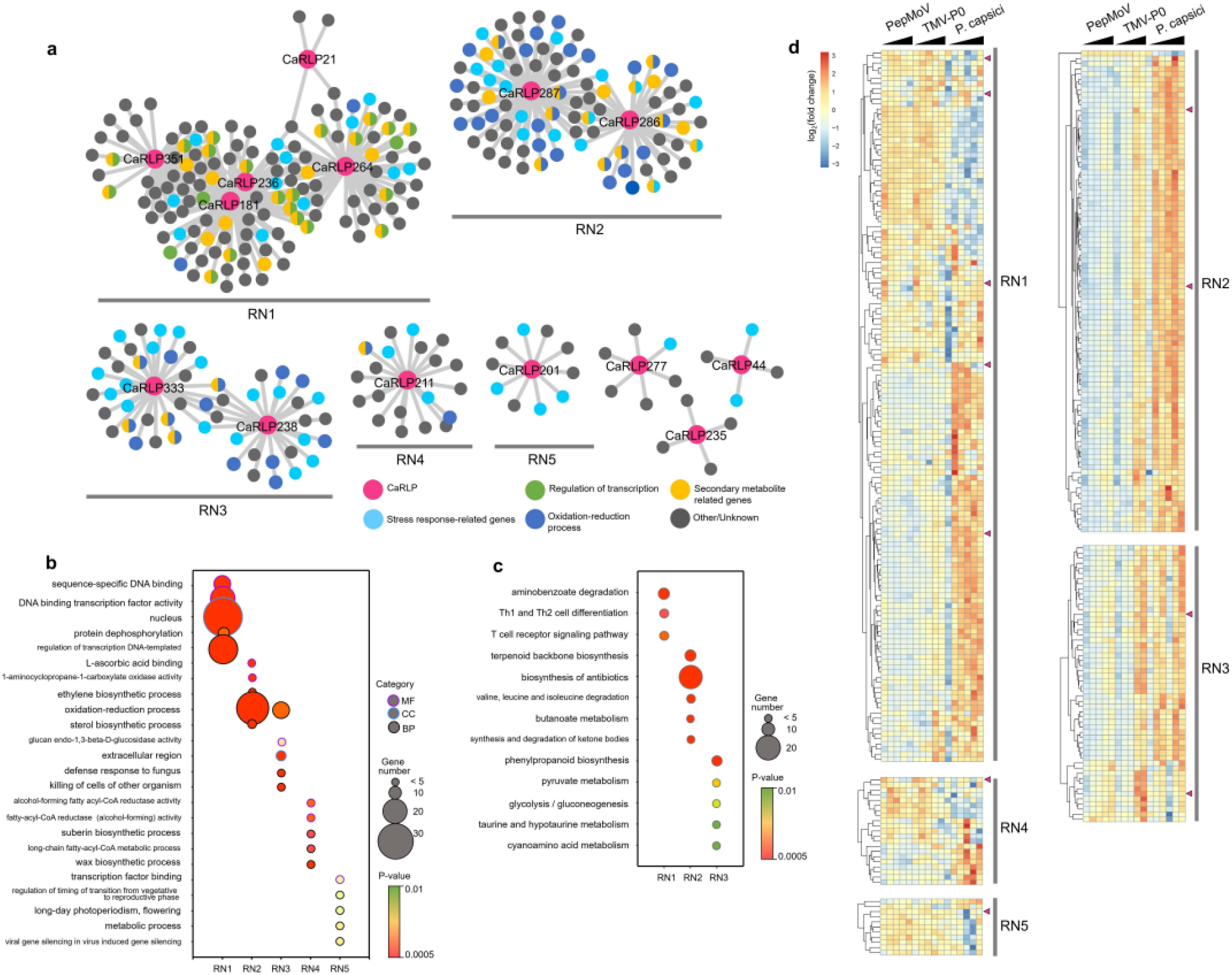
Identification of biotic stress-responsive core *CaRLPs* in a universal GCN. (a) Intersection of three GCNs of PepMoV-, TMV-P0- or *P. capsici*-infected plants. The co-expression network modules containing more than ten nodes were designated as RN1–RN5. Magenta nodes represent *CaRLPs*, and other colored nodes represent their annotated functions by GO analysis. (b) Top 5 GO categories enriched (*P* < 0.01) in five modules (RN1 to RN5). The Y-axis and X-axis in the bubble plot represent the GO category and different modules, respectively. The purple, blue, and black borders of the circle represent molecular function (MF), cellular component (CC), and biological process (BP), respectively. The size and color of each bubble represent the number of DEGs and *P*-value for each category, respectively. (c) Bubble plot showing the results of KEGG pathway enriched analysis of five co-expression modules (RN1 to RN5). The Y-axis and X-axis in the bubble plot represent the enriched KEGG pathways and network modules, respectively. The size and color of each bubble represent the number of DEGs and *P*-value for each category, respectively. (d) Expression profiles of genes in RN1–RN5 modules. Expression values were normalized relative to the value of control samples (Mock). Module names are indicated on the right hand side of the heatmap. Magenta triangles on the right side of the heat map indicate hub *CaRLPs* in each module. Black triangles on the top of the heat map represent the time-course of each pathogen infection.

Next, we performed GO and KEGG enrichment analyses and expression analysis to determine the biological processes and pathways affected during the plant immune responses of *CaRLPs* and associated pepper genes in RN. Thus, we focused on GO terms belonging to the BP category in these modules. Stress-related genes were enriched in RN modules (Fig. 4a and 4b). Notably, genes involved in stress response were higher enriched in RN2 and RN3 modules. We focused on GO terms belonging to the BP category in these modules. GO terms related plant defense mechanisms such as “oxidation-reduction process,” “sterol biosynthetic process” and “ethylene biosynthetic process” were highly enriched in the RN2 module, and “oxidation-reduction process,” “defense response to fungus,” and “cellular oxidant detoxification” were highly enriched in RN3. As mentioned above, oxidation-reduction and cellular oxidant detoxification occur in plants in response to pathogen attack. Most pathogenic fungi and oomycetes uptake sterols from the external environment, most likely from the host cell membrane, during pathogenesis^50^. In addition, ethylene acts as a signaling molecule during stress^51^. The results of KEGG pathway analysis revealed the enrichment of defense related pathways such as “biosynthesis of antibiotics,” “terpenoid backbone biosynthesis,” and “phenylpropanoid biosynthesis” in the RN (Fig. 4c). “Terpenoid backbone biosynthesis” and “phenylpropanoid biosynthesis” pathways produce secondary metabolites, which are involved in plant defense^52^. Taken together, these findings support that genes co-expressed with *CaRLPs* in the RN are involved in biotic stress response. We also examined the expression profiles of genes in the RN (Fig. 4d). Based on expression patterns, genes in RN1 were divided into two types: those highly up-regulated in response to both PepMoV and TMV-P0, and those up-regulated mainly in response to *P. capsici*. Genes in RN2 and RN3 modules were up-regulated in response to *P. capsici*. The expression of genes in each module was significantly correlated, indicating that these genes were tightly connected each other. Thus, these results suggest that hub *CaRLPs* in the RN play a role in resistance to multiple biotic stresses.

### Functional validation of core *CaRLPs* involved in HR-like response to pathogen invasion

We hypothesized that core *CaRLPs*, i.e., hub genes in universal GCN, are involved in resistance response to biotic stresses. To decipher the core *CaRLPs* of the GCN, which potentially function in biotic stress response, we performed loss-of-function analysis of nine *CaRLP* genes; each of these genes was silenced by virus-induced gene silencing (VIGS) in the pepper cultivar ‘Nockwang.’ These nine genes included 2 genes not belonging to the RN (*CaRLP35* and *71*) and 7 core *CaRLPs* belonging to the RN (*CaRLP181*, *211*, *264*, *277*, *286*, *287* and *351*); the remaining 7 of the 14 core *CaRLPs* were excluded from this analysis, as their nucleotide sequence was none-specific for VIGS assay. VIGS constructs were constructed by cloning a sequence unique to each of the nine *CaRLPs* into a *Tobacco rattle virus* (TRV) vector; notably, because *CaRLP286* and *287* exhibit high level of sequence similarity, both these genes could be silenced using a single construct containing a sequence common to the two genes. The expression level of each *CaRLP* was significantly lower in *CaRLP*-silenced pepper plants than in the *TRV2-GFP* control (Supplementary Fig. S5), although no significant phenotypic difference was observed between *TRV2-CaRLP* and control plants (Supplementary Fig. S6), indicating that the eight *CaRLP* constructs did not affect the growth and development of pepper plants.

To investigate whether the silencing of *CaRLPs* affects HR, a form of programmed cell death (PCD), upon TMV-P0 infection, we simultaneously inoculated *CaRLP*-silenced pepper plants and control plants with TMV-P0, and monitored their phenotypes. The number of HR lesions on TMV-P0-inoculated leaves was significantly lower in *TRV2-CaRLP264*, *−277*, *−286/287* and *−351* lines than in *TRV2-GFP* plants at 48 hpi (Fig. 5a). The level of HR in these four *CaRLP*-silenced lines was decreased by 0.22–0.72-fold compare with that in control plants (Fig. 5b). By contrast, the silencing of other *CaRLP* genes did not cause any significant change in the number of HR lesions. These results suggest that these *CaRLP* genes are involved in the activation of defense mechanisms and PCD upon pathogen infection in pepper.

**Fig. 5.**
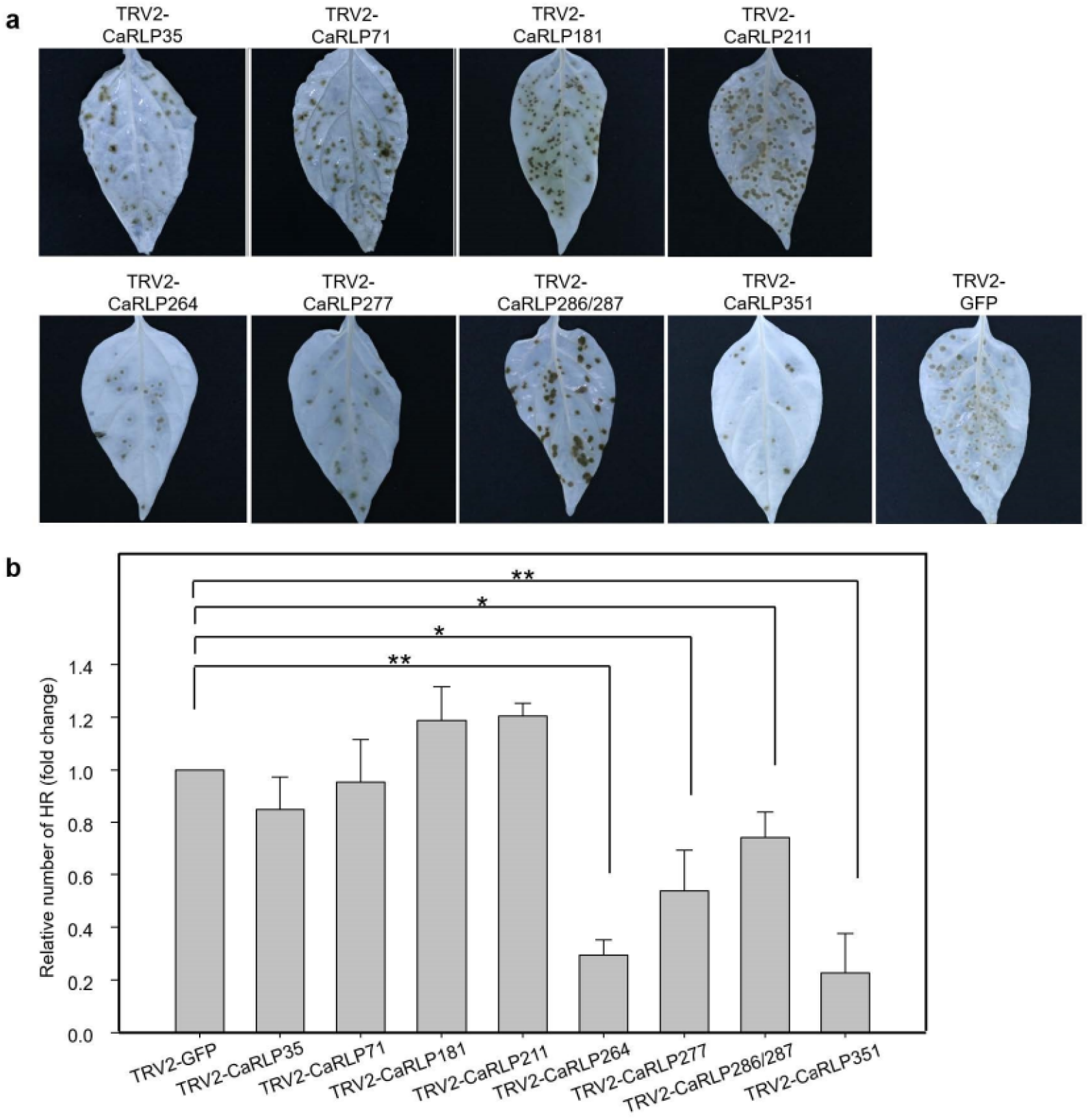
Assessment of HR lesions in CaRLPs-silenced peppers to TMV-P0 infection. (a) Photographs showing TMV-P0-inoculated leaves of *CaRLP*-silenced plants. Photos were taken at 3 dpi. Chlorophyll was removed using ethyl alcohol. (b) Reduced HR lesion numbers in *CaRLP*-silenced plants inoculated with TMV-P0. Data indicate mean ± standard error (SE) of three independent experiments (*n* = 24). Asterisks indicate statistically significant differences compared with the *TRV2:GFP* control (\**p* < 0.05, \*\**p* < 0.01; Student’s *t*-test).

### Enhanced defense responses of core *CaRLP*-silenced pepper plants to various pathogens

To determine whether the core *CaRLPs* are involved in broad-spectrum resistance to various pathogens, we tested the response of *CaRLP*-silenced and *TRV2-GFP* control plants to *Xanthomonas axonopodis* pv. *glycines* 8ra (Xag8ra), *Ralstonia solanaceaerum* (Rsol) and *P. capsici*. We examined three different responses of *CaRLP*-silenced to the above mentioned pathogens; for instance, non-host resistance to Xag8ra, host resistance to Rsol and susceptible response to *P. capsici*. We selected three *CaRLPs* (*CaRLP264*, *277* and *351*), which showed the most significant difference in the resistance response to TMV-P0 infection (Fig. 5).

To investigate the role of *CaRLPs* during HR response of non-host resistance^53,54^, control plants (*TRV2-GFP*) and *CaRLP*-silenced plants (*TRV2-CaRLP264*, - *CaRLP277* and - *CaRLP351*) were infiltrated with Xag8ra (10^8^ cfu/ml) (Fig. 6a). Xag8ra-inoculated plants of each *CaRLP*-silenced line showed significantly reduced HR-like cell death compared with control plants at 48 hpi. In addition, quantification of ion leakage from the inoculation-induced lesion showed that conductivity of each *CaRLP*-silenced line was approximately 1.5–2-fold lower than that of control plants (Fig. 6b). This suggests that the core *CaRLPs* play a crucial role in HR-based immunity of pepper plants against Xag8ra.

**Fig. 6.**
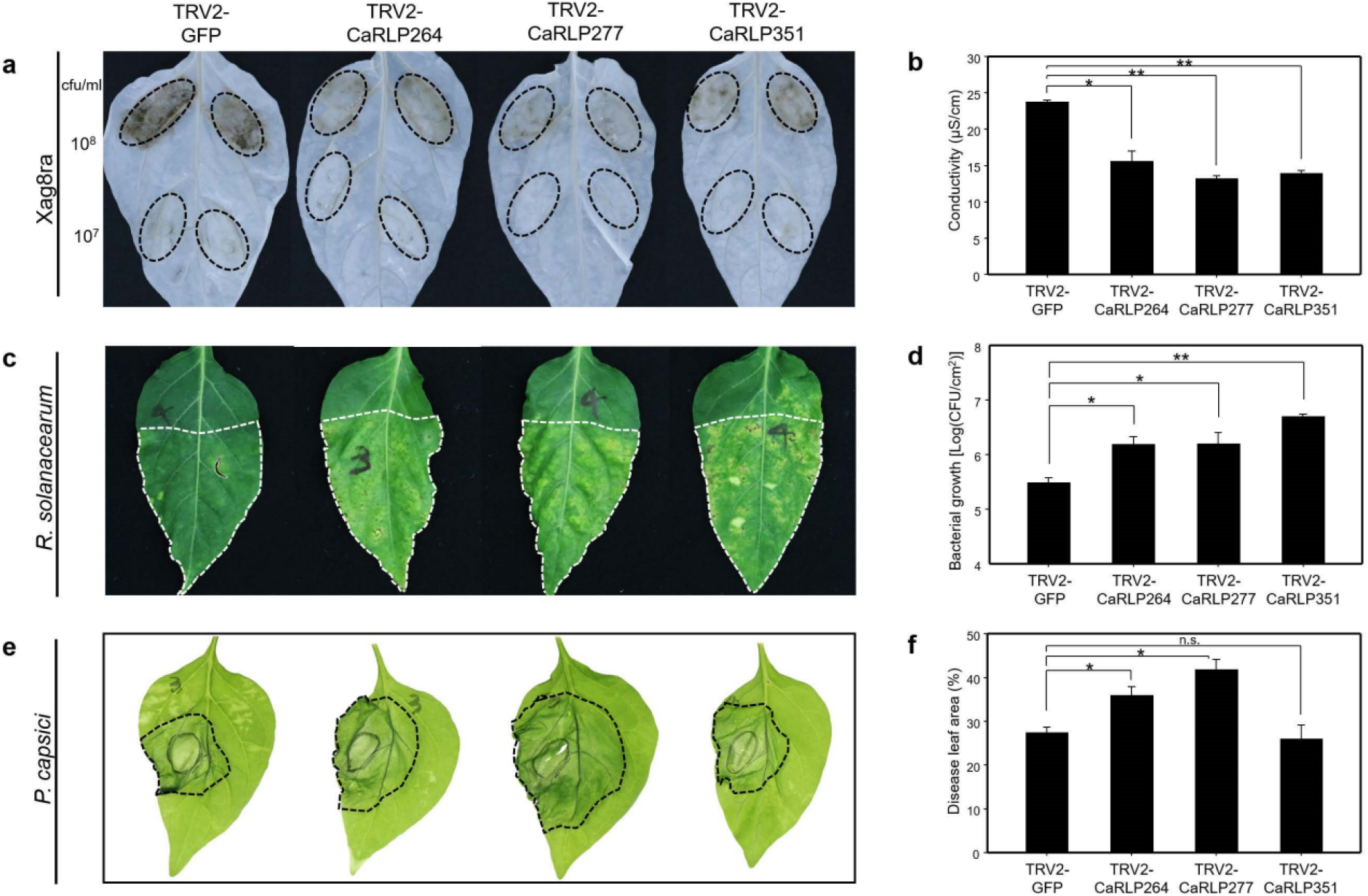
Broad-spectrum resistance of core *CaRLP*-silenced peppers against various pathogens. (a) Response to Xag8ra inoculation on leaves of *CaRLP*-silenced plants. Photos were taken at 2 dpi. Chlorophyll was removed using ethyl alcohol. (b) Ion leakage data of Xag8ra-inoculated leaves. Ion leakage was measured using leaf disks from inoculation lesion of 10^8^ cfu/ml of Xag8ra. (c) Response of *R. solanacearum* inoculation on leaves of *CaRLP*-silenced plants. Photos were taken at 8 dpi. (d) Bacterial growth of *R. solanacearum* in silenced plants. (e) Disease symptoms caused by *P. capsici* inoculation of the leaves of *CaRLP*-silenced plants and control plants. Photos were taken at 3 dpi. (f) Disease lesion width normalized relative to the total leaf area. All data indicate mean ± SE from three independent experiments. Asterisks indicate statistically significant differences compared with the *TRV2:GFP* control (\**p* < 0.05, \*\**p* < 0.01; Student’s *t*-test). n.s., not significant (*p* > 0.05).

Next, we performed leaf infiltration of Rsol, the causal agent of bacterial wilt disease, into *CaRLP*-silenced plants of *C. annuum* cultivar ‘MC4,’ which is resistant to Rsol^55^. Three *CaRLP*-silenced plants rapidly developed leaf wilting symptoms including necrosis and yellowing, as observed in susceptible plants but not in control plants (Fig. 6c). Furthermore, the growth of Rsol was increased significantly by approximately 5–15-fold in *CaRLP*-silenced plants compared with that in control plants at 5 dpi (Fig. 6d). These findings suggest that silencing *CaRLP264*, *CaRLP277*, and *CaRLP351* enhances the susceptibility to Rsol.

Finally, the leaves of CaRLP-silenced and control plants were also challenged with *P. capsici*, and disease development was examined at 3 dpi. The *CaRLP264*- and *CaRLP277*-silenced plants showed larger disease lesions than control plants, whereas *CaRLP351*-silenced plants showed no significant difference in lesion size compared with the control (Fig. 6e and 6f). This suggests that CaRLP263 and CaRLP277 are involved in the defense response to *P. capsici*. Taken together, plants of each *CaRLP*-silenced line consistently showed significant suppression of broad-spectrum defense against plant pathogens including viruses, bacteria, and oomycetes. Overall, our data suggest that core *CaRLPs* of the universal GCN perform conserved functions and confer resistance against multiple biotic stresses. Thus, these *CaRLPs* could be used to engineer cultivars with broad-spectrum resistance against diverse pathogens.

## DISCUSSION

Plants sense pathogens via both cell surface and intracellular receptors. RLPs represent the primary layer of defense against pathogen infection in the innate immune system. In the present study, we identified a large number of *CaRLP* genes in the pepper genome, and selected variable biotic stress-responsive *CaRLP* genes as components of GCNs using 102 RNA-seq datasets. We demonstrated that three hub *CaRLPs* in the universal GCN confer broad-spectrum resistance against diverse pathogens.

A large percentage of genes in eukaryotic genomes are organized in clusters of various sizes and gene densities. Clusters containing resistance gene analogs (RGAs) including NLR, RLKs and RLPs have been reported in plants^24,44,56^. In tomato and pepper, several genes related to RGAs are localized in clusters on various chromosomes^56–58^. Consistent with this data, we observed that *CaRLPs* belonging to the same phylogenetic group were mostly located in the same cluster, and thus showed uneven chromosomal distribution (Fig. 1). Information on the chromosomal location of *CaRLPs* would be highly valuable for the identification of functional RGAs.

GCNs could provide important clues for the characterization of novel genes, based on the analysis of potentially functionally associated co-expressed genes, using large-scale gene expression datasets^28^. Here, we attempted to infer the function of *CaRLPs* under various biotic stresses by the analysis of GCNs derived from a large number of RNA-seq datasets of *C. annuum* ‘CM334.’ GCNs were constructed using *CaRLPs* as hub genes in two steps: construction of large CaRLP-GCNs, based on the RNA-seq of each biotic stress, and construction of the intersection of CaRLP-GCNs, according to the type of biotic stress. Four large CaRLP-GCNs were constructed, each corresponding to four biotic stresses and containing 1,073–10,878 genes. These CaRLP-GCNs showed that *CaRLPs* are co-expressed with numerous other pepper genes under various biotic stresses. This result is consistent with previous studies, which showed that plant response to pathogens is extensively regulated at the transcriptional level^59–61^.

The intersection of these four GCNs led to the construction of RN (Fig. 3 and Fig. 4). The RNA-seq data in this study was obtained from *C. annuum* cultivar ‘CM334,’ which is resistant to TMV-P0, PepMoV, and *P. capsici* but susceptible to TMV-P2. Consequently, GO enrichment analysis of genes in the RN revealed the enrichment of various stress related terms such as “oxidation-reduction process,” “defense response to fungus,” “response to biotic stimulus,” and “cellular response to oxidative stress” (Fig. 4). In addition, numerous genes were enriched not only in stress related GO terms but also in transcription regulation (Fig. 4). Most of genes in the RN enriched under “regulation of transcription” encoded transcription factors, such as WRKY, AP2/ERF domain-containing proteins. These transcription factors play critical roles in abiotic and biotic stresses^62^. For example, WRKY proteins are involved in RLP-mediated defense response. Signal transduction of AtRLP51 was mediated by BDA1, an Ankyrin-repeat-containing protein with four transmembrane domains, to provoke plant defense response through the activation of WRKY70^63,64^. Taken together, these data suggest that *CaRLPs* in universal GCNs could be co-regulated with transcription factors under biotic stress.

We hypothesized that core *CaRLPs* in the universal GCN (RN) are involved in the response to biotic stresses. To test our hypothesis, we characterized the function of core *CaRLPs* using VIGS. We were not able to develop stable transgenic plants in pepper because of the limitation of transgenic system for pepper and the low regeneration rate of pepper plants under in vitro conditions^65^. Of the six *CaRLPs* tested in this study, plants silenced for the expression of three *CaRLPs* showed reduced HR-like cell death upon TMV-P0 and Xag8ra inoculation (Fig. 5 and Fig. 6). By contrast, the silencing of each *CaRLP* significantly enhanced the disease susceptibility of pepper plants to *Rsol* and *P. capsici* compared with control plants (Fig. 6). These data suggest that the core *CaRLPs* in the universal GCN perform conserved functions and induce broad-spectrum resistance against plant pathogens. Thus, construction of a universal GCN from comprehensive transcriptome datasets could provide useful clues for uncovering the roles of genes in various biological processes.

Resistance gene-mediated immunity is highly effective immune systems against specific pathogens. On the other hands, PRRs, located on the plant cell surface, could confer resistance to a broad range of pathogens. In previous studies, few plant PRRs showed broad-spectrum resistance to pathogens. Expression of the Arabidopsis elongation factor Tu (EF-Tu), one of the PRRs, in *N. benthamiana* and tomato increased resistance to *Pseudomonas*, *Agrobacterium*, *Xanthomonas* and *Ralstonia*^4^. In potato, the elicitin response (ELR) receptor-like protein associates with the immune co-receptor BAK/SERK3, and mediates broad-spectrum recognition of elicitin proteins form several *Phytophthora* species^5^. In addition, suppression of the pepper lectin receptor kinase gene *CaLecRK-S.5*, which acts as a PRR, showed enhanced susceptibility to PepMoV, *Xanthomonas*, and, *P. capsici* ^66^. In this study, through generation of a conserved GCN, we identified PRRs involved in broad-spectrum resistance against diverse plant pathogens. Three *CaRLPs* (*CaRLP264*, *277* and *351*) enhanced susceptibility to TMV-P0, *Xanthomonas*, *Ralstonia*, and *P. capsici* (Fig. 5 and Fig. 6). Thus, these *CaRLPs* could potentially be used to develop Solanaceae crop cultivars with broad-spectrum resistance against diverse pathogens. Overall, a universal GCN with comprehensive RNA-seq datasets could provide key insights to unveil gene functions in biological processes.

## MATERIALS AND METHODS

### Identification of *CaRLP* genes

A total of 13 characterized plant *RLP* genes (Supplementary Table S8) were used to obtain *CaRLP* gene sequences, which were used to build an hidden Markov model (HMM) domain with the HMMER software package (version 3.0; http://hmmer.org/), and identified putative RLP-encoding genes against the *C. annuum* ‘CM334’ v. 1.55 genome. Then, tBLASTn searches were performed using the HMMER domain from amino acid sequences encoded by the pepper genome (threshold: 10^−4^). Consequently, 600–750 hits to genes in the pepper genome were obtained from the BLAST output, accounting for 7,376 genes in total. This gene set was processed to remove redundant sequences by manual curation, thus obtaining 784 non-redundant candidate genes. The structure of CaRLPs was annotated using Pfam^67^ and SMART ^68^ databases, and genes with kinase and NB-ARC domains were filtered out using Pfam IDs PF07714.12, PF00069.20 and PF00931, respectively. Finally, 438 CaRLPs from the ‘CM334’ genome.

### Phylogenetic analysis and classification of CaRLPs

The CaRLPs were classified based on the results of phylogenetic analysis and sequence similarity-based clustering, as described previously^24,44^. Clustering analysis of full-length amino acid sequences of Arabidopsis, tomato, pepper and reported RLPs was performed by OrthoMCL^69^. RLPs within the same cluster were determined to be identical subgroups to the phylogenetic subgroups. RLPs clustered as singletons (mostly partial and short sequences) were identified using a BLASTP search against identified RLPs, and subgroups were assigned. We designated the known RLP names according to the corresponding pepper RLP groups.

A conserved domain of an HMM profile was built based on the amino acid sequence of conserved C3-D region^25^ of known RLPs. A phylogenetic tree was constructed based on the C3-D domain of 56 AtRLPs^22,25^, 176 SlRLPs^24^, 438 CaRLPs, and 13 RLPs reported by HMM search (E-value < 0.001). *RLP* genes containing less than 80% of the full-length C3-D domain sequence were excluded. Multiple sequence alignment of the C3-D domains of RLPs was performed using MUSCLE (http://www.ebi.ac.uk/Tools/msa/muscle/). The alignment result was used to build a phylogenetic tree using PhyML (http://www.phylogeny.fr/), with default parameters (SH-like approximate likelihood-ratio test for branch support), and the resulting phylogenetic trees were edited using the MEGA8 software (http://www.megasoftware.net/).

### Chromosomal location, physical cluster, and motif analyses

The chromosomal location of *CaRLPs* was determined based on the genome sequence of the pepper cultivar ‘CM334’^42^. MapChart^70^ was used to draw the location of RLPs on chromosomes. Physical clustering of *CaRLPs* in the pepper genome was determined based on two criteria: 1) the gene cluster spans a region of 200 kb or less; and 2) the cluster contains less than eight non-*RLP* genes between two *CaRLPs*^24,44^.

Conserved motifs in CaRLPs were identified using the MEME suite (http://meme-suite.org/tools/meme), with default settings except for the following parameters: maximum number of motifs, 20; minimum width of motifs, 15; maximum width of motifs, 200. Subsequently, MAST (http://meme-suite.org/tools/mast) was carried out on datasets including protein sequences of CaRLPs and known RLPs with default E-values.

### RNA-seq library construction

Changes in the expression profiles of *CaRLPs* upon *P. capsici* infection were investigated in the pepper cultivar ‘CM334’ by RNA-seq analysis. Leaves of 4–5-week-old pepper plants were infiltrated with *P. capsici* (5 × 10^4^ zoospore/ml), and infected leaves were collected at 0, 1, 2, 4, 6, 12, and 24 hpi in three biological replicates. Total RNA was isolated using the TRIzol Reagent (Invitrogen, Carlsbad, CA, USA), according to the manufacturer’s instructions. RNA-seq libraries were constructed as described previously^45^. All 39 RNA-seq libraries (21 libraries of *P. capsici*-infected samples, and 18 libraries of control samples) were sequenced using Illumina HiSeq 2000 (Illumina Inc., San Diego, CA, USA).

### RNA-seq data analysis

Quality control of RNA-seq data of *P. capsici*-infected samples was performed by removing low-quality reads and possible contaminants, as described previously^40,41^. Adapter and low-quality sequences were filtered using Cutadapt^71^ and Trimmomatic^72^, based on the Phred quality threshold of 20. In addition, transcriptome data of pepper plants infected by TMV-P0, TMV-P2, and PepMoV were obtained from previous studies^43,45^ to analyze the expression of *CaRLPs*.

### Gene expression analysis

The expression profiles of *CaRLPs* under biotic stresses were analyzed using RNA-seq data of TMV-P0-, TMV-P2-and PepMoV-inoculated pepper plants and that of *P. capsici*-inoculated pepper plants. Sequence reads from all RNA-seq datasets were aligned to the ‘CM334’ reference genome using Hisat2^73^. Filtered clean reads of virus RNA-seq and *P. capsici* RNA-seq were normalized into reads per kilobase per million mapped reads and fragments per kilobase per million mapped fragments, respectively. DEGs were identified using the DESeq2 package (FDR < 0.05)^74^. The expression patterns of DEGs were visualized using ComplexHeatmap^75^.

### GCN construction and GO and KEGG enrichment analyses

The GCN was constructed from 102 RNA-seq datasets using the exp2net function of the mlDNA package^76^, and inferred using the Pearson’s product moment correlation coefficient at a significance level of *P* < 0.01. In next step, genes co-expressed with *CaRLPs* were identified by filtering the correlation coefficient (|r| > 0.8) and only directed interaction. The GCN was visualized using Cytoscape v3.4.0 ^77^. To identify gene networks involved in different stress responses, GCNs containing *CaRLP* genes were extracted by different combinations of all stresses using Merge Tools in Cytoscpae. GO and KEGG enrichment analyses were performed by GOseq^78^ in R packages using Pepper v1.55 genome annotation from BLAST2GO^79^.

### VIGS

Pepper cultivars *C. annuum* ‘Nockwang’ and ‘MC4’ were used to analyze the effect of *CaRLP* gene silencing on defense response. Seedlings with two fully expanded cotyledons were for the VIGS assay. The 3’ or 5’ untranslated region (UTR) of eight *CaRLP* genes (*CaRLP35*, *CaRLP71*, *CaRLP181*, *CaRLP211*, *CaRLP264*, *CaRLP277*, *CaRLP286*/*287* and *CaRLP351*) was amplified and cloned into the pTRV2 vector. The resulting pTRV2-CaRLP constructs were transformed into *Agrobacterium tumefaciens* strain GV3101. VIGS was conducted as described previously^80^. Plants infiltrated with *pTRV2-GFP* or *pTRV2-PDS* with pTRV1 were used as a control. One leaf was harvested from each *CaRLP*-silenced plant for RNA extraction and the measurement of silencing efficiency.

### Pathogen inoculation

All pathogen inoculations were performed on the 3^rd^ and 4^th^ true leaves of *CaRLP*-silenced and control pepper plants at 4–5 weeks after the VIGS assay. Plants were challenged with three different types of pathogens (including viruses (TMV-P0, Xag8ra), bacteria (Rsol) and oomycete (*P. capsici*). The TMV-P0 inoculum was prepared from 1 g of infected *N. benthamiana* leaves using 10 ml of 0.1 M phosphate buffer (pH 7.0). TMV-inoculated leaves were monitored and harvested at 3 days post-inoculation (dpi). To assess the formation of lesions on TMV-inoculated leaves, chlorophyll was removed using ethyl alcohol. To conduct the Rsol-response assay, Rsol *‘*SL1931’ was cultured first in TZC agar medium at 28°C for 2 days and then in CPG medium at 28°C for 24 h, and then suspended in distilled sterile water. The Rsol suspension was diluted to a concentration of 10^5^ cfu/ml, and infiltrated into the leaves of *CaRLP*-silenced pepper plants. Subsequently, Rsol-inoculated plants were grown in a growth chamber at 28 ± 2°C, 70% relative humidity and 16h-light/8h-dark photoperiod, and inoculated leaves were harvested at 5 dpi. Inocula of Xag8ra and *P. capsici* were prepared as described previously ^53,80^. The Xag8ra culture was suspended in 10 mM MgCl_2_, and then diluted to 10^7^ and 10^8^ cfu/ml concentrations. The Xag8ra-inoculated leaves were harvested at 2 dpi and used for measuring conductivity and detecting cell death. To prepare the *P. capsici* inoculum, the released zoospores were collected and diluted in distilled sterile water to a concentration of 1 × 10^5^ spores/ml. The *P. capsici* suspension was infiltrated into the leaves of *CaRLP*-silenced pepper plants, and harvested at 3 dpi. All pathogen inoculation were conducted in at least three independent experiments, with 8–12 plants for per experiment.

### Quantification of ion leakage

Ion leakage from Xag8ra-inoculated leaves was measured as described previously^53^. Sixteen leaf disks (each 1 cm in diameter) were excised from 4–6 plants of each *CaRLP*-silenced line, and floated on 15 ml of sterile distilled water for 2 h at room temperature. Then, the electrolyte leakage from leaf discs was measured by a conductivity meter (Eutech con 510; Thermo scientific, Waltham, MA, USA).

### Bacterial cell counting

In Rsol-inoculated pepper plants, bacterial cell growth was measured in planta, as described previously^55^, with slight modifications. Six leaf disks (each 1 cm in diameter) were excised from Rsol-inoculated leaves of 3–4 plants of each *CaRLP*-silenced line at 5dpi. Leaf disks were ground in sterile distilled water, and serial dilutions were plated on CPG agar medium supplemented with gentamycin. The plates were incubated at 28°C, and bacterial cells were counted after 2 days.

## ACKNOWLEDGEMENTS

This research was supported by the National Research Foundation of Korea (NRF) funded by the Korean Government (NRF-2017R1E1A1A01072843 and 2019R1C1C1007472). We appreciate the support from the KRIBB initiative program.

## AUTHOR CONTRIBUTIONS

WHK, BP, and JSK collected samples and performed experiments. YMK and JL generated RNA-seq data. NK and WHK analyzed transcriptome. WHK and SIY conceived and designed the experiments, organized and wrote the manuscript, and supervised the project. All authors read and approved the final manuscript.

