## Supplementary information for "Universal gene co-expression network reveals receptor-like protein genes conferring broad-spectrum resistance in pepper (*Capsicum annuum* L.)"

**Table S2. Physical clusters of *RLP* genes in the pepper (*C. annuum*) genome**

| Chromosome | No. of clusters | Average total no. of genes in cluster | Mean cluster size (kb) | No. of singletons |
|---|---|---|---|---|
| 1 | 3 (9)[a] | 3.0 | 109.9 | 9 |
| 2 | 5 (13) | 2.6 | 41.1 | 15 |
| 3 | 2 (7) | 3.5 | 8.6 | 9 |
| 4 | 12 (64) | 5.3 | 288.6 | 17 |
| 5 | 6 (29) | 4.8 | 110.8 | 6 |
| 6 | 4 (8) | 2.0 | 6.4 | 11 |
| 7 | 3 (6) | 2.0 | 9.3 | 14 |
| 8 | 7 (32) | 4.6 | 53.8 | 6 |
| 9 | 3 (7) | 2.3 | 36.9 | 7 |
| 10 | - | - | - | 8 |
| 11 | - | - | - | 6 |
| 12 | 9 (52) | 5.8 | 127.1 | 15 |

[a] Numbers of genes in clusters are given in parentheses.

**Supplementary Table S3. Identification of conserved motifs in *CaRLP*s.**

| Motif | Width | Multilevel consensus sequence[a] | Encoding domain[b] |
|---|---|---|---|
| 1 | 34 | HNKFSGEIPQQMANLTFLAVLDLSHNHLTGCIPQ | LRR |
| 2 | 21 | LQVLDLSHNHFTGTIPQCIGN | LRR |
| 3 | 21 | TYIDLSHNKFEGHIPEEIGDL | LRR |
| 4 | 41 | HGNKLTGKIPRSICNCKYLQVLDLGDNHLTGTIPMWLGNL P | LRR |
| 5 | 41 | NLQYLDLSHNQFTGPIPSWIGNLKNLQYLDLSHNQFTGEI P | LRR |
| 6 | 29 | GKQFHTFENSSYEGNDGLCGFPLSKDCGG | LRR-like |
| 7 | 21 | LRVLDLSHNKFTGHIPASMGN | LRR |
| 8 | 21 | DCCSWDGVTCDEMTGHVIELD | LRR-like |
| 9 | 29 | FWKAAAMGYGCGLCIGLAIIYIMASTHYM | Transmembrane region |
| 10 | 41 | QVLSLRSNKFHGPIRTSRTENMFPQLRMIDLSCNAFTGNL P | LRR |
| 11 | 20 | LEVLDMSHNNFSGTIPTWFC | LRR |
| 12 | 29 | GEIPEEIGNLTNLEYLDLSYNQFTGPIPS | LRR |
| 13 | 21 | YYQDSVAVVTKGLEMEVVRIL | |
| 14 | 29 | SQLQGKFHSNSSLFQLHHLQRLDLSYNDF | LRR |
| 15 | 21 | HFCHEDQATALLQWKAMFTDQ | LRR |
| 16 | 21 | QLTGTIPEEIGNLKNLQYLDL | |
| 17 | 21 | KLQYLDLRNNKLHGPIPDWIW | LRR |
| 18 | 21 | YLDMSHNNFSGEIPASICNLT | LRR |
| 19 | 29 | WLARIIEELEHKIMMRRRKKQRRQRNYRR | |
| 20 | 21 | FCNLKQLEYLDLSYNHFSGEI | LRR |

[a] Consensus sequences of conserved motifs were obtained from the analysis of the 438 *CaRLP*s and well-known 12 *RLP* genes with MEME.

[b] Determination with SMART by default value

**Supplementary Table S4. Statistics of the transcriptome data of *Phytophthora capsici*-infected plants of pepper cultivar 'CM334.'**

| | Parameters | Time point (hours post infection) | | | | | | |
|---|---|---|---|---|---|---|---|---|
| | | 0hr | 1hr | 2hr | 4hr | 6hr | 12hr | 24hr |
| *P. capsici* infected CM334 | Total reads† | - | 37,137,516 | 40,790,294 | 44,779,996 | 42,701,370 | 32,401,138 | 21,325,716 |
| | Clean reads‡ | - | 26,728,028 | 29,702,402 | 32,457,810 | 31,064,928 | 27,502,404 | 20,319,004 |
| | Mapped genome§ | - | 21,054,267 | 24,845,907 | 27,868,854 | 26,875,857 | 23,564,408 | 17,249,912 |
| | Mapped genome (%)¶ | - | 78.73 | 83.66 | 85.86 | 86.52 | 85.65 | 84.90 |
| | Mapped CDS†† | - | 14,243,216 | 17,052,885 | 20,136,003 | 19,526,251 | 16,203,755 | 11,630,852 |
| | Mapped CDS (%)‡‡ | - | 53.27 | 57.39 | 62.04 | 62.86 | 58.67 | 57.11 |
| Mock | Total reads† | 35,565,322 | 23,199,976 | 22,876,380 | 21,196,338 | 9,335,632 | 9,525,190 | 15,925,524 |
| | Clean reads‡ | 25,488,950 | 22,276,968 | 21,952,974 | 20,318,436 | 8,952,528 | 9,121,060 | 14,962,112 |
| | Mapped genome§ | 19,918,135 | 16,759,438 | 17,433,131 | 16,704,454 | 7,466,855 | 7,620,179 | 12,151,980 |
| | Mapped genome (%)¶ | 78.15 | 75.24 | 79.41 | 82.22 | 83.41 | 83.47 | 81.17 |
| | Mapped CDS†† | 13,304,093 | 11,214,602 | 11,819,497 | 11,351,964 | 4,971,644 | 4,917,850 | 7,952,918 |
| | Mapped CDS (%)‡‡ | 52.20 | 50.35 | 53.84 | 55.86 | 55.55 | 53.81 | 53.08 |

† Results of the sum of three replicates.

‡ Trimmed reads with the sum of three replicates.

§Number of reads (sum of three replicates) mapped to the 'CM334' genome.

¶ Percent mapped reads (average of three replicates).

†† Number of reads (sum of three replicates) mapped to the coding sequences (CDSs) of 'CM334.'

‡‡ Percent reads mapped to the CDSs of 'CM334' (average of three replicates).

**Supplementary Table S5. Statistical summary of RNA-seq data used in this study.**

| Pathogen | Treatment | Time point | Read type | Processed data (Gb) [†] | References |
|---|---|---|---|---|---|
| Virus | Mock | 0, 0.5, 4, 24, 48, 72h | SE [‡] | 12.00 | (Kim et al., 2018) |
| | PepMoV | 0.5, 4, 24, 48, 72h | SE | 8.49 | (Kim et al., 2018) |
| | TMV-P0 | 0.5, 4, 24, 48, 72h | SE | 5.68 | (Kim et al., 2018) |
| | TMV-P2 | 0.5, 4, 24, 48, 72h | SE | 0.82 | (Kang et al., 2016) |
| Oomycete | Mock | 0, 1, 2, 4, 6, 12, 24h | PE [§] | 17.21 | In this study |
| | *P. capsici* | 1, 2, 4, 6, 12, 24h | PE | 22.88 | In this study |

[†] Total number of clean seqeunces of three replicates at each time point

[‡] Single end

[§] Paired end

**Supplementary Table S8. Information of known plant *RLP*s used in this study**.

| Name | Species | Function | References |
|---|---|---|---|
| *Cf-2* | Tomato | *Cladosporium fulvum* resistance | (Dixon et al., 1996) |
| *Cf-4* | Tomato | *Cladosporium fulvum* resistance | (Thomas et al., 1997) |
| *Cf-5* | Tomato | *Cladosporium fulvum* resistance | (Dixon et al., 1998) |
| *Cf-9* | Tomato | *Cladosporium fulvum* resistance | (Jones et al., 1994) |
| *Cf-4A* | Tomato | *Cladosporium fulvum* resistance | (Takken et al., 1998) |
| *Ve1* | Tomato | *Verticillium* resistance | (Kawchuk et al., 2001) |
| *Ve2* | Tomato | Interaction with SOBIR1 | (Kawchuk et al., 2001) |
| *LeEIX2* | Tomato | Xylanase elicitor perception of *Trichoderma viride* | (Ron and Avni, 2004) |
| *ReMAX* | Arabidopsis | Recognition of eMAX from Xanthomanads | (Jehle et al., 2013) |
| *RFO2* | Arabidopsis | *Fusarium oxysporum* resistance | (Shen and Diener, 2013) |
| *HcrVf2* | Apple | *Venturia inaequalis* resistance | (Vinatzer et al., 2001) |
| *TMM* | Arabidopsis | Stomatal ditribution | (Yang and Sack, 1995) |
| *CLV2* | Arabidopsis | Meristem development | (Taguchi-Shiobara et al., 2001) |

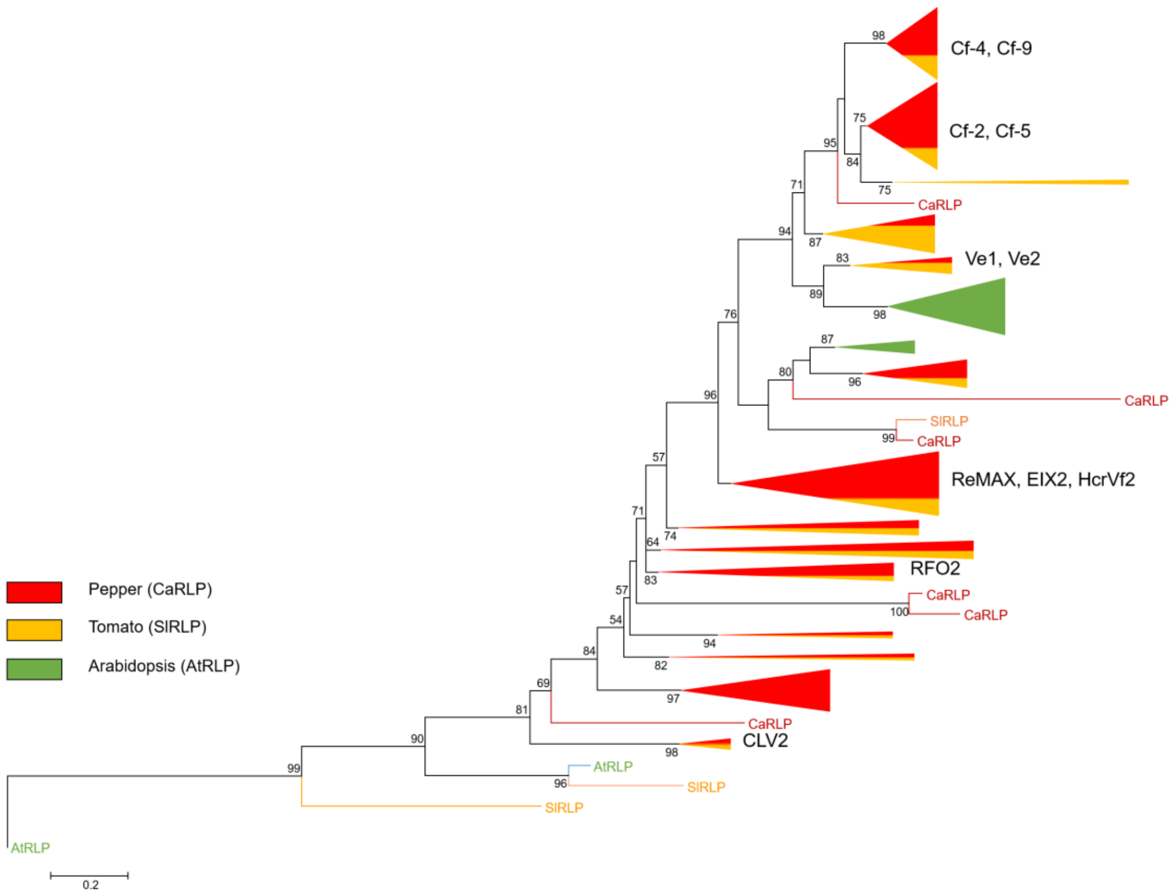


**Supplementary Figure S1. Phylogenetic relationships of pepper, tomato, and Arabidopsis *RLP* genes.** The alignment and phylogenetic analysis was performed with C3-D domains. The phylogenetic tree was constructed using the Maximum-likelihood method in PhyML. *RLP*s of pepper, tomato, and Arabidopsis are represented by red, yellow, and green, respectively. Bootstrap values over 50 are indicated above branches. The clades including known *RLP* genes are indicated by *RLP* gene names on the right side of the clade.

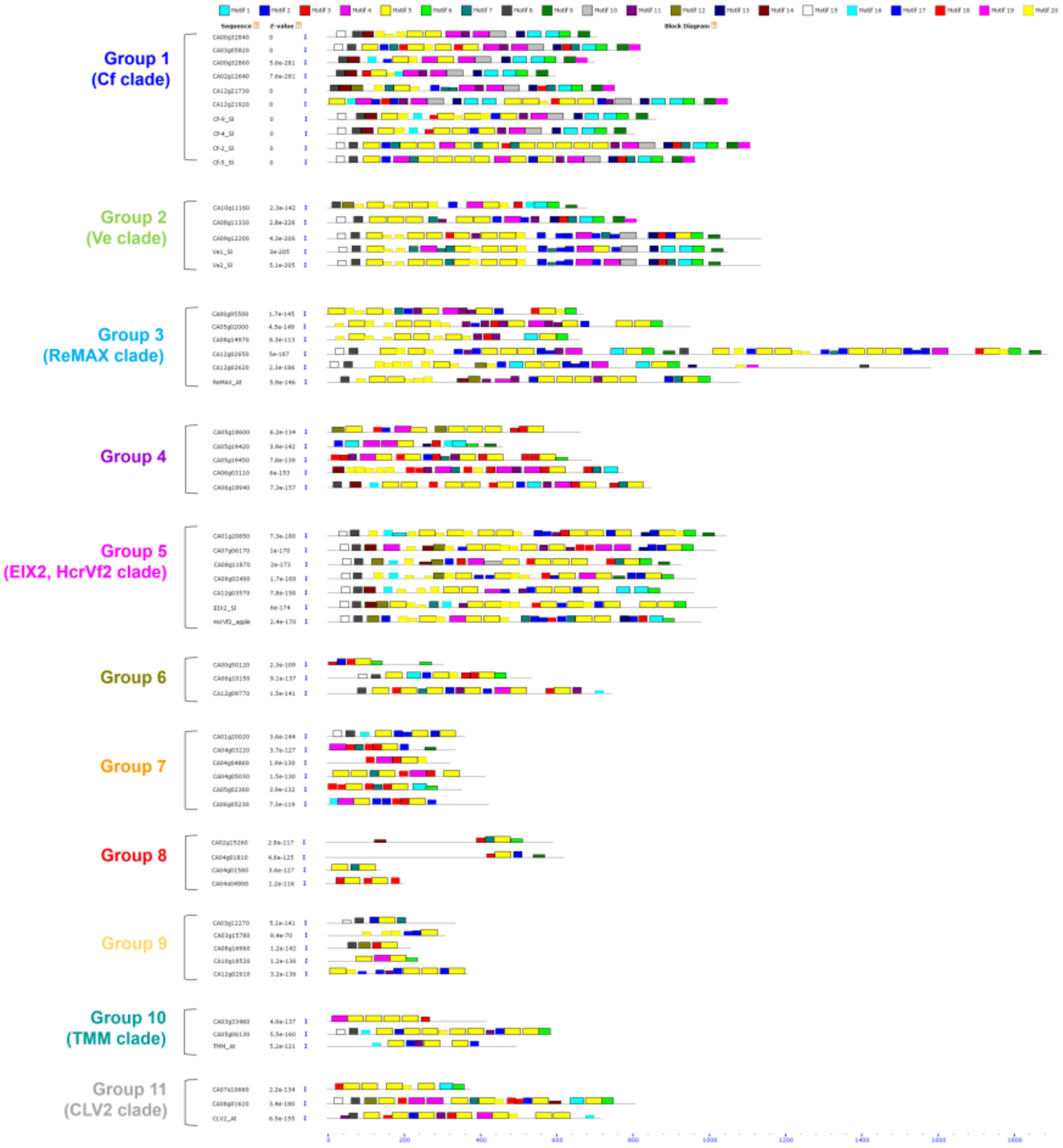


**Supplementary Figure S2. Identification of *RLP* motifs in pepper.** The 20 motifs were identified by MEME. The different colored boxes indicate different motifs and their potion in each *RLP* protein. Several subgroups were distinguished by phylogenetic analysis. The left side of figure indicate clade name according to phylogenetic tree.

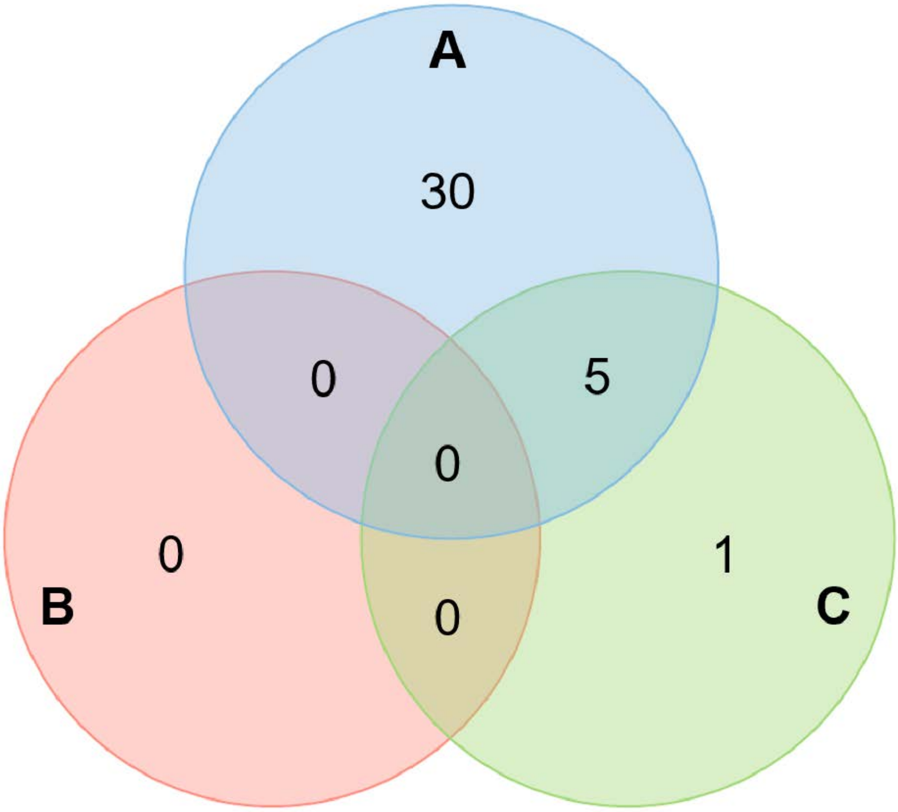


**Supplementary Figure S3. Venn diagram for CaRLP-DEGs from three different virus treated RNA-seq.** The number in each circle represents the number of CaRLP-DEGs of TMV-P0 (A), TMV-P2 (B), and PepMoV (C) treated RNA-seq.

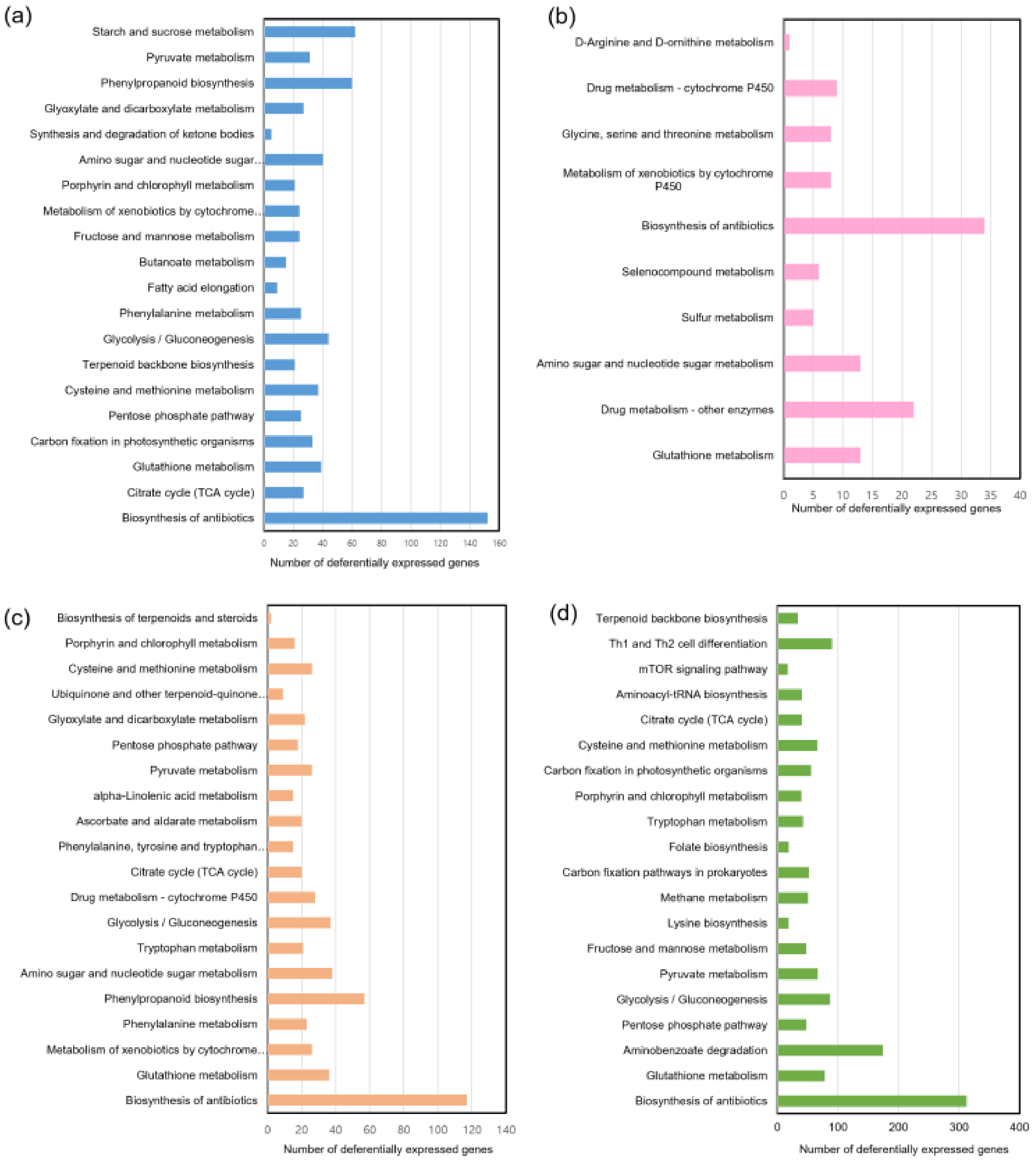


**Supplementary Figure S4. KEGG pathway analysis of four CaRLP-GCNs.** Top 20 KEGG pathways were selected from TMV-P0 (a), PepMoV (c), and P. capsici (d)- CaRLP-GCN. All annotated pathways were selected from KEGG analysis of TMV-P2 –CaRLP-GCN in figure (b).

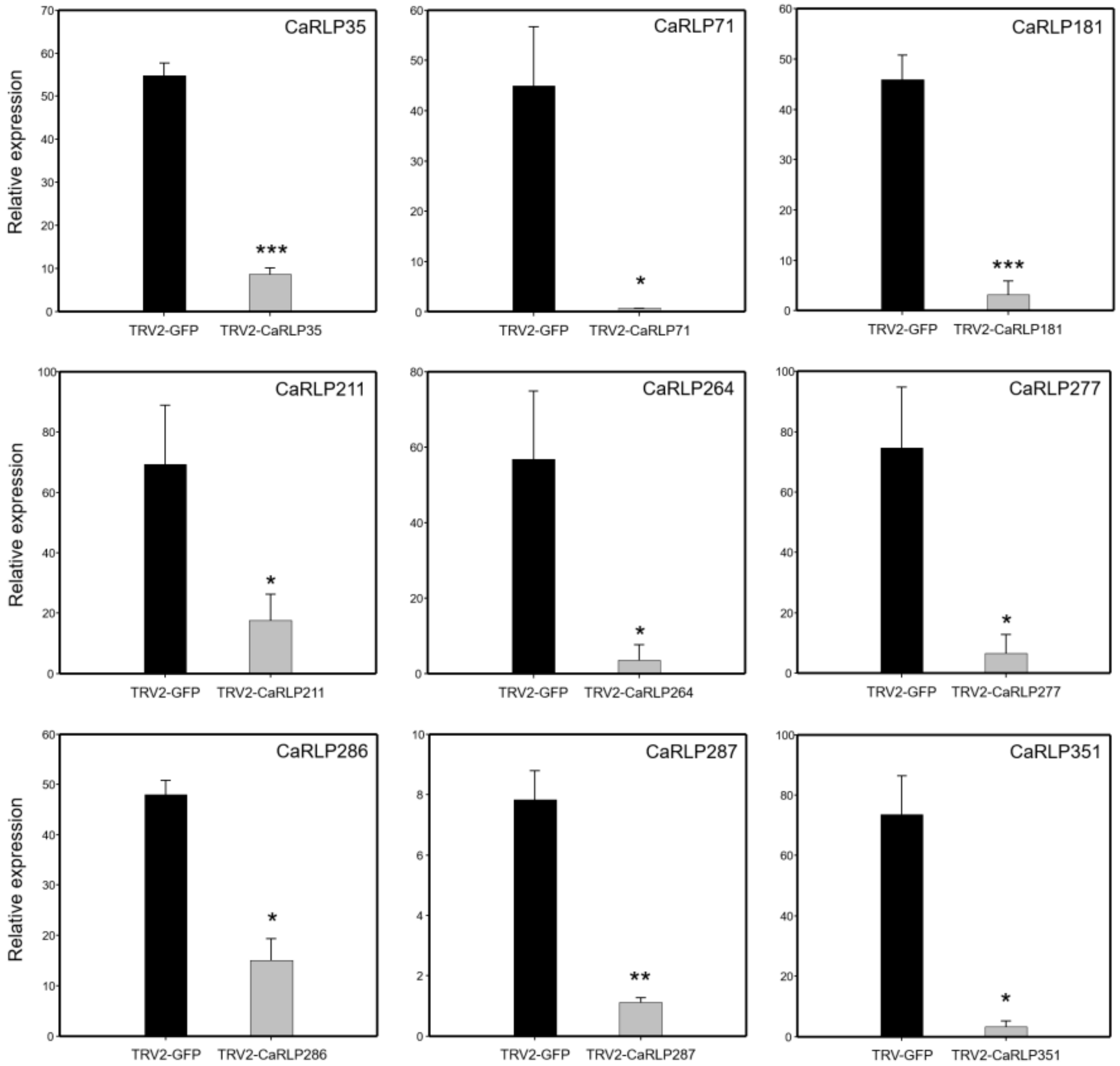


**Supplementary Figure S5. Reverse transcription-polymerase chain reaction (RT-PCR) analyses of the each CaRLP in silenced pepper leaves.** TRV2-GFP was used as a control. Expression values were normalized to levels of *CaActin* gene expression. Data represent means±SD from three independent experiments (Student's *t*-test, *$p$<0.05, **$p$<0.01, ***$p$<0.005).

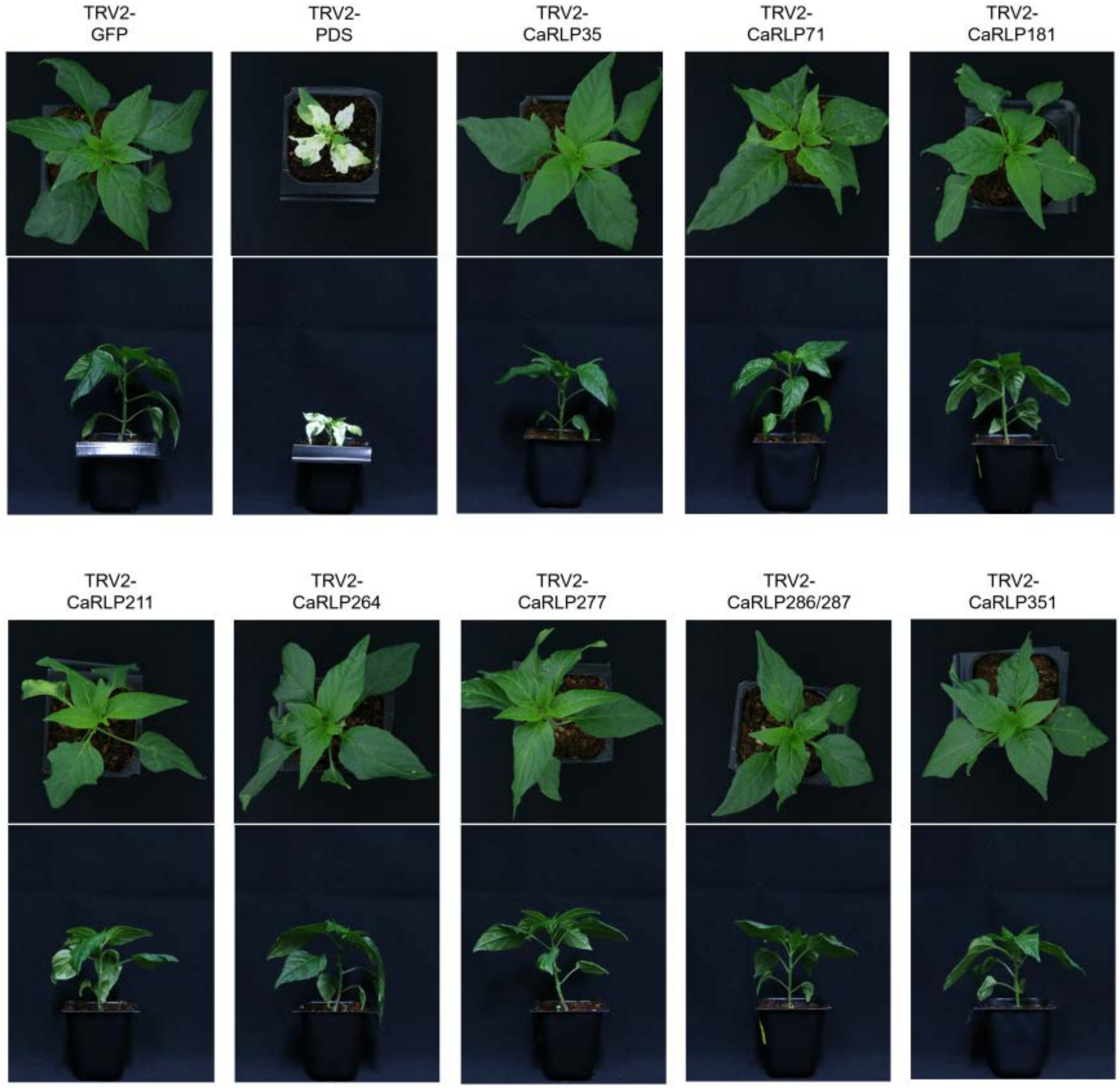


**Supplementary Figure S6. Silencing of CaRLP in pepper plant.** Pepper seedlings were inoculated with Agrobacterium containing the TRV2-CaRLP plasmid by syringe infiltration. Photograph was taken 28 days after inoculation.
